# Voluntary co-contraction of ankle muscles alters motor unit discharge characteristics and reduces estimates of persistent inward currents

**DOI:** 10.1101/2024.02.28.582534

**Authors:** Matheus M. Gomes, Sophia T. Jenz, James A. Beauchamp, Francesco Negro, C.J. Heckman, Gregory E.P. Pearcey

## Abstract

Motoneuronal persistent inward currents (PICs) are both facilitated by neuromodulatory inputs and highly sensitive to local inhibitory circuits (e.g., Ia reciprocal inhibition). Methods aimed to increase group Ia reciprocal inhibition from the antagonistic muscle have been successful in decreasing PICs, and the diffuse actions of neuromodulators released during activation of remote muscles have increased PICs. However, it remains unknown how motoneurons function in the presence of simultaneous excitatory and inhibitory commands. To probe this topic, we investigated motor unit (MU) discharge patterns and estimated PICs during voluntary co-contraction of ankle muscles, which simultaneously demands the contraction of agonist-antagonist pairs. Twenty young adults randomly performed triangular ramps (10s up and down) of both co-contraction (simultaneous dorsiflexion and plantarflexion) and isometric dorsiflexion to a peak of 30% of their maximum muscle activity from a maximal voluntary contraction. Motor unit spike trains were decomposed from high-density surface electromyography recorded over the tibialis anterior (TA) using blind source separation algorithms. Voluntary co-contraction altered motor unit discharge rate characteristics, decreasing estimates of PICs by 20% (4.47 pulses per second (pps) vs 5.57 pps during isometric dorsiflexion). These findings suggest that, during voluntary co-contraction, the inhibitory input from the antagonist muscle overcomes the additional excitatory and neuromodulatory drive that may occur due to the co-contraction of the antagonist muscle, which constrains PIC behavior.

**KEY POINTS:** Voluntary co-contraction is a unique motor behavior that concurrently provides increases in excitatory and inhibitory inputs to motoneurons.

During co-contraction of agonist-antagonist pairs, agonist motor unit discharge characteristics are altered, consistent with reductions in persistent inward current magnitude.

Reciprocal inhibition from the antagonist likely becomes proportional to the increase in neural drive to the agonist, dampening the magnitude of persistent inward currents.

## INTRODUCTION

It is now well-established that monoaminergic inputs from the brainstem facilitate persistent inward currents (PICs), which modulate the excitability of the motor pool and are necessary for normal motor behavior (Heckman *et al*., 2005; Heckman & Enoka, 2012; Johnson *et al*., 2017). PICs are depolarizing currents generated by dendritic voltage sensitive ion channels that increase motoneuronal excitability by amplifying and prolonging synaptic inputs (Heckman *et al*., 2005). The magnitude of PICs depends directly on the level of monoamines released, in particular norepinephrine (NE) and serotonin (5HT) (Lee & Heckman, 2000; Heckman *et al*., 2005; Heckman & Enoka, 2012), but also the amount and pattern of inhibition from local circuits (Heckman *et al*., 2009; Binder *et al*., 2020).

The majority of evidence on the inhibitory control of PICs arises from animal studies and computer models (Hultborn *et al*., 2003; Kuo *et al*., 2003; Hyngstrom *et al*., 2007; Bui *et al*., 2008) but the magnitude of PICs in human motoneurons can be estimated quite easily due to recent advances in technology and analysis techniques (Klotz *et al*., 2023; Möck & Del Vecchio, 2023). There remains a scarcity of research in humans, however, that establishes a relationship between reciprocal inhibition and PIC magnitude (Thorstensen, 2022). Mesquita et. al. (2022) recently found that stimulation of the common peroneal nerve, which induces reciprocal inhibition of the plantar flexors, can reduce PIC magnitude estimated from motor units decomposed from the human gastrocnemius and we have shown that vibration of an antagonist muscle tendon can reduce estimates of PIC magnitude, which we believe were due to reciprocal inhibition triggered by the vibratory stimuli of the antagonist muscles (Pearcey *et al*., 2022). Since vibration is a crude method of inducing reciprocal inhibition, as it can induce several types of sensory input (e.g., non-locally mechanoreceptors, heteronomous muscle spindles) which may also impact motoneuron excitability, we remained interested in exploring further experimental paradigms that exemplify the control of PIC magnitude.

Examining the behavior of motor units of the agonist muscle during the voluntary muscle contraction of its antagonist (i.e., voluntary co-contraction) is a promising way to deepen understanding about the interplay between inhibitory control and neuromodulatory mechanisms underlying motor unit discharge characteristics. Contrary to most motor actions, where contraction of an agonist muscle causes reciprocal inhibition of its antagonistic pair, an isometric co-contraction task demands simultaneous contraction of antagonistic muscles and likely alters the structure of excitatory, inhibitory, and neuromodulatory commands to motoneurons.

Therefore, to understand the effect of voluntary muscle co-contraction on motor unit properties in humans, we compared motor unit discharge rate profiles during both isometric dorsiflexion and co-contraction tasks about the human ankle. Since PICs are highly sensitive to inhibition and voluntary co-contraction of the antagonist will likely impart reciprocal inhibition onto the agonist muscle, we hypothesized that co-contraction would alter motor unit discharge rate profiles, which would indicate lower PIC magnitude.

## METHODS

### Participants and Ethical Approval

Twenty healthy adults were recruited for this study. However, four participants were excluded because they were not able to perform the co-contraction task properly. Data from sixteen adults (7 Female; 30.3 ± 6.2 years) were analyzed. Participants had no experience with co-contraction training and no history of neuromuscular, musculoskeletal, or other abnormalities that would prevent them from completing the tasks described below. All participants provided written informed consent, approved by the Institutional Review Board of Northwestern University (STU00202964, STU00216332).

### Experimental Protocol

Participants first performed three maximum voluntary isometric contractions (MVCs) of five seconds for each condition (i.e., dorsiflexion, plantarflexion, and co-contraction), with 90-second rest intervals between each attempt. If there was greater than 10% difference between MVC trials in peak torque (during dorsiflexion and plantarflexion) or rectified TA/SOL EMG amplitude (during co-contraction), additional trials were added. Verbal encouragement was given to participants during MVC trials to ensure maximal muscle contraction. The maximum torque and EMG amplitudes achieved during MVCs were used for subsequent normalization of all trials.

Participants next performed a familiarization comprised of 10 to 20 triangular-shaped ramps of TA contractions and co-contractions. The triangular ramp consisted of 10s linear ascending and descending phases to a peak of 30% of the rectified TA EMG achieved during maximal voluntary co-contraction. Participants received real-time visual feedback of the TA EMG during all submaximal ramp contractions on a large monitor that was positioned 2m away (Figure 1).

**Figure 1.**
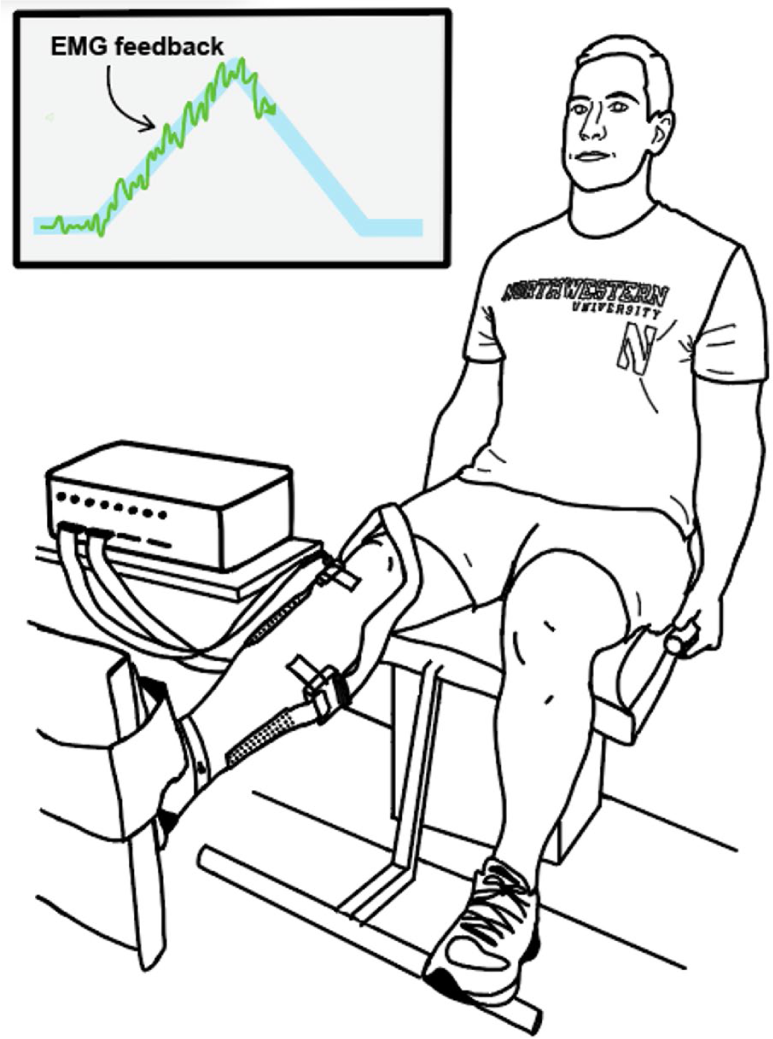
– Participant performing a co-contraction triangular ramp. The blue triangular trace represents the contraction intensity (target) to be performed and the green line represents a smoothed TA electromyogram visual feedback.

For the experimental protocol, participants performed four co-contraction ramps and four dorsiflexion ramps in a randomized order. Since co-contraction demands simultaneous activation of the plantar flexor and dorsiflexor muscles, we expected very little ankle torque. As such, if participants performed more than 20% of their maximum torque in either direction (dorsiflexion and plantarflexion) during co-contraction, additional trials were added as needed.

### Torque recording

Isometric torque during dorsiflexion, plantarflexion, and co-contraction were collected using a Systems 2 isokinetic dynamometer (Biodex Medical System, Shirley, NY). Participants were seated with their right foot in a footplate attachment, the hip in approximately 100° of flexion, knee at approximately 170°, and ankle at 100° (Figure 1). Velcro straps were wrapped around the foot to secure it to the plate and prevent movement. Torque was sampled at 2048 Hz and smoothed offline with a 10 Hz low-pass filter (fifth-order Butterworth filter).

### High-density surface electromyography (HD-sEMG) Recording

Before electrode placement, excess hair was removed and the skin overlying the muscle of interest was lightly abraded. High-density surface EMG (HD-sEMG) electrodes (64 electrodes, 13×5, 8mm I.E.D., GR08MM1305, OT Bioelettronica, Inc., Turin, IT) were placed longitudinally over the muscle belly of the tibialis anterior (TA) and soleus (SOL) (Rainoldi *et al*., 2004), which are dorsiflexor and plantar flexor muscles of the ankle, respectively (Figure 1). A reference electrode strap was placed around the right ankle, overlying the medial and lateral malleolus. The HD-sEMG signals were collected in differential mode, sampled at 2048 Hz, amplified x150, and band-pass filtered at 10-500Hz using a Quattrocento signal amplifier (OT Bioelettronica, Inc., Turin, IT). EMG and torque were temporally synced with a 1-second TTL pulse transmitted to the Quattrocento at the onset of each trial. In order to provide EMG feedback to participants, the signal from one channel of the TA electrode was amplified (x16) using OTbio+ software from the analog output feature. The EMG channel was chosen based on a central location on the electrode and signal to noise quality. This EMG channel was output to a NI DAQ (NI-USB-6218) where a custom MATLAB script was used to process and display the feedback onto the screen. The feedback EMG was lowpass filtered (1.8 Hz), the resting baseline was removed, and averaged over 60 ms intervals before being provided back to the participant.

### Data Analysis

#### Torque

From the MVC trials, the maximum voluntary torque (MVT) achieved was used for subsequent normalization of all trials. Regarding the co-contraction ramp trials, only trials in which the participant did not achieve significant torque in any direction (i.e., less than 20% of the maximum dorsiflexion and plantarflexion torque) were included in subsequent analyses.

#### Motor unit decomposition

All surface HD-sEMG signals were initially bandpass filtered at 10–500 Hz (second-order, Butterworth) and visually inspected to remove substantial artifacts or noise. Individual motor unit spike trains were decomposed from the remaining HDsEMG channels using a convolutive blind-source separation algorithm with a silhouette threshold of 0.87 (Negro *et al*., 2016). After decomposition, motor unit spike trains were manually edited by a trained technician. This inspection used a custom-made graphical user interface in MATLAB to correct minor errors made by the decomposition algorithm using well-validated local re-optimization methods to improve motor unit spike trains similar to the techniques used in recent studies(Boccia *et al*., 2019; Afsharipour *et al*., 2020; Del Vecchio *et al*., 2020; Martinez-Valdes *et al*., 2020; Jenz *et al*., 2023). Instantaneous discharge rates of each motor unit spike train were determined by computing the inverse of the interspike interval and smoothed using support vector regression (Beauchamp *et al*., 2022) with custom-written MATLAB scripts. Within these scripts, the initial, peak, and final discharge rates were extracted from the smoothed spike trains. Ascending duration was calculated as the time that a motor unit exhibited sustained discharge before peak torque, and descending duration as the time a motor unit exhibited sustained discharge from peak torque to derecruitment. Finally, recruitment threshold was calculated as the level of torque at the first motor unit spike.

#### Estimates of Persistent Inward Currents (PICs)

Effects of PICs on motor unit discharge patterns can be estimated by quantifying the amount of onset-offset hysteresis (i.e., ΔFrequency, ΔF) of a higher-threshold (test) motor unit with respect to the discharge rate of a lower-threshold (reporter) unit (Gorassini et al. 1998, 2002). Rather than providing ΔF values for each test-reporter unit pair, we calculated ‘unit-wise’ values, which gives each test unit one ΔF value based on the average values obtained from multiple reporter units. Criteria for inclusion of ΔF values from motor unit pairs were that 1) the test unit was recruited at least 1 second after the reporter unit to ensure full activation of the PIC (Bennett *et al*., 2001; Powers *et al*., 2008; Hassan *et al*., 2020), 2) test unit-reporter unit pair exhibited rate-rate correlations ≥ 0.7 to ensure motor unit pairs received common synaptic drive (Gorassini *et al*., 2004; Udina *et al*., 2010; Stephenson & Maluf, 2011; Vandenberk & Kalmar, 2014), and 3) the reporter unit displayed a discharge range of at least 0.5 pps while the test unit was active (Stephenson & Maluf, 2011).

#### Geometric Analysis

We used an additional method, referred to as brace height (Beauchamp et al., 2023) to quantify the nonlinearity of motor unit discharge rate with respect to EMG output. Using the smoothed discharge rate of a single motor unit, we quantified the maximum orthogonal deviation between this smoothed discharge and an expected linear increase from motor unit recruitment to peak discharge, with the maximal deviation representing the “brace height”. Brace height values were normalized to the height of a right triangle with a hypotenuse from recruitment to peak MU discharge. Motor units with a negative acceleration phase slope, brace height exceeding 200% after normalization, or peak discharge occurring after peak force were excluded. To distinguish between the secondary and tertiary phases of motor unit discharge, we calculated slopes for the initial acceleration phase and subsequent attenuation phase of motor unit discharge, along with the angle formed between these phases (Beauchamp et al., 2023).

#### Tracking Motor Units

Motor units from isometric and co-contraction trials were tracked using a custom MATLAB script, which tracked motor units using blind source separation filters of the motor unit spike trains (Francic & Holobar, 2021; Oliveira & Negro, 2021; Goodlich *et al*., 2023). The dataset of tracked motor units were analyzed using the same methods as the full dataset, and results are reported.

### Statistical analysis

Statistical analyses were performed using R Statistical Software (v4.1.0; R Core Team 2021). To determine if the fixed effect of contraction type predicted variables of interest, we used linear mixed effects models (lmer R package; v1.1.27.1; (Bates *et al*., 2015). To determine significance, we applied Satterthwaite’s method for degrees of freedom (lmerTest R package; v3.1.3; (Kuznetsova *et al*., 2017). Estimated marginal means and effect size (Cohen’s *d*) were computed for each variable (emmeans R package, v1.8.0,(Lenth RV, Bolker B, Buerkner P, Gine-Vázquez I, Herve M & Love J, Miguez F, Riebl H, 2023). Results are reported as emmeans ± standard deviation. Effect size was used to determine the effect of condition from the estimated marginal means of isometric and co-contraction data from the model. All data were visualized in R (ggplot R package, v3.3.6) (Wickham H, Chang W, Henry L, Pedersen TL, Takahashi K & Woo K, Yutani H, Dunnington D, 2023)

## RESULTS

### EMG and Torque During Isometric and Co-contraction

To evaluate how effectively participants performed the challenging co-contraction task, we compared net ankle torque and EMGs in both the agonist and antagonist between the isometric and co-contraction protocols. The peak dorsiflexion torque was much higher in the isometric condition (43.1 ± 2.90 %MVT) than during co-contraction (14.0 ± 2.91 %MVT χ^2^_(1)_ = 162.57, *P* < 0.001, *d* = 3.67). Peak agonist (TA) EMG did not differ during isometric (26.0 ± 1.13 %MVEMG) or co-contraction (26.9 ± 1.14 %MVEMG; χ^2^_(1)_ = 1.1634, *P* = 0.281, *d* = −0.20). However, the peak EMG of the antagonist muscle (SOL) was much lower during isometric (6.7 ± 1.62 %MVEMG) than co-contraction (12.2 ± 1.64 %MVEMG; χ^2^_(1)_ = 17.15, *P* < 0.001, *d* = −0.803). The similarity in TA EMG magnitude during both contractions allows for the comparison of motor units between the two conditions.

### Motor units decomposed (per trial and participant for each condition) and recruitment torques

Across all the trials, 1,031 units were decomposed from the isometric condition and 996 units were decomposed from the co-contraction condition. Participants had more difficulty performing co-contraction successfully, and therefore there were fewer valid attempts (trials) at co-contraction. On average 16.60 units were decomposed per trial for the isometric condition, and 17.47 units per trial for the co-contraction condition. Figure 3 shows the distribution of motor unit recruitment was similar in isometric and co-contraction.

**Figure 2.**
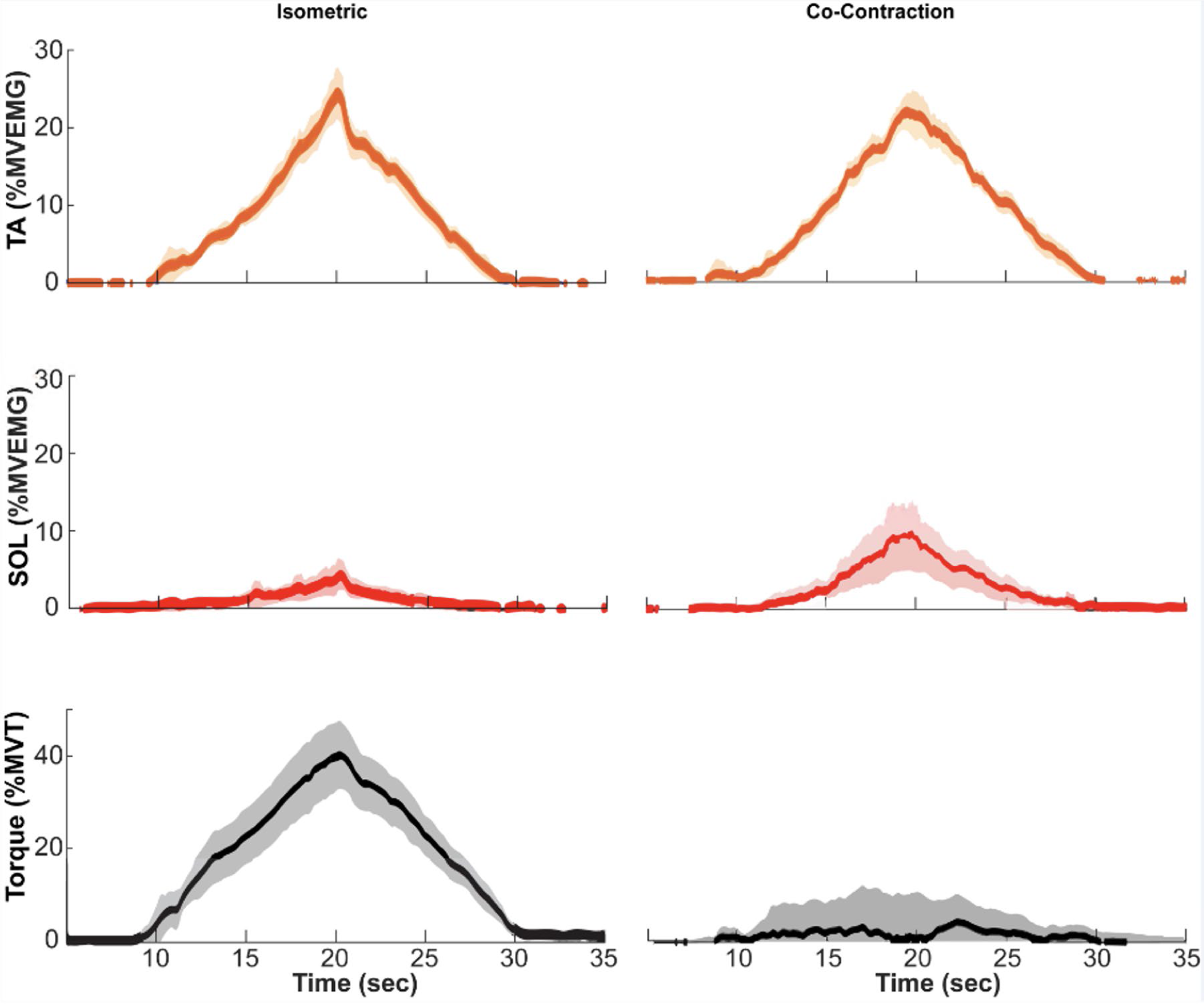
– Normalized EMG torque and dorsiflexion torque during isometric and co-contraction shows participants correctly performed co-contraction. Normalized EMG for the agonist muscle, TA, is shown in orange and antagonist, SOL, in red. Normalized dorsiflexion torque is shown in black. Abbreviations: MVEMG; maximal voluntary EMG; MVT; maximal voluntary torque.

**Figure 3.**
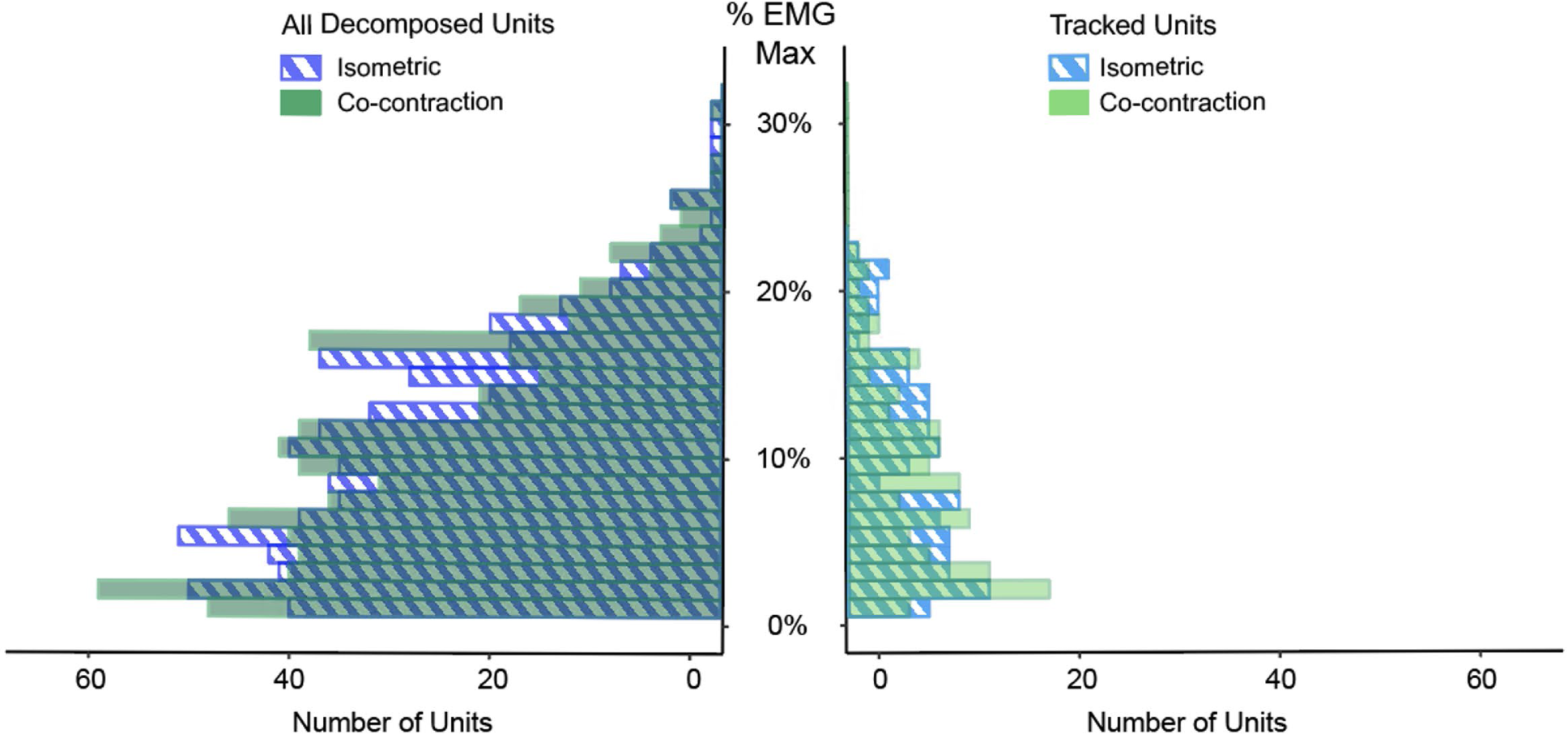
– The number of motor units decomposed by %EMG recruited for the full and tracked datasets for isometric (blue hatched) and co-contraction (solid green) conditions.

### Motor Unit Discharge Rate Differs between Conditions

To gain insight into motor unit activity during the triangular ramps we quantified discharge rate at recruitment, derecruitment, and the peak discharge rate (Table 1). In the full dataset, the motor unit discharge rate at recruitment was higher during the isometric condition (χ^2^ = 15.135, *P* < 0.001, *d* = 0.22), but the discharge rate at derecruitment was lower in isometric (χ^2^ = 4.1794, *P* = 0.041, *d = −0.115*). Peak discharge rate was higher during co-contraction (χ^2^ = 10.16, *P* = 0.001, *d* = −0.19). For tracked MUs, discharge rate was similar at recruitment (χ^2^ = 1.374, *P* < 0.241, *d* = 0.142) and peak (χ^2^ = 2.122, *P* = 0.145, *d =* −0.177), but was lower in isometric at derecruitment (χ^2^ = 4.805, *P* < 0.028, *d* = −0.268). These values show motor unit discharge of the agonist muscle is altered when the antagonist is simultaneously active (Figure 4).

**Figure 4.**
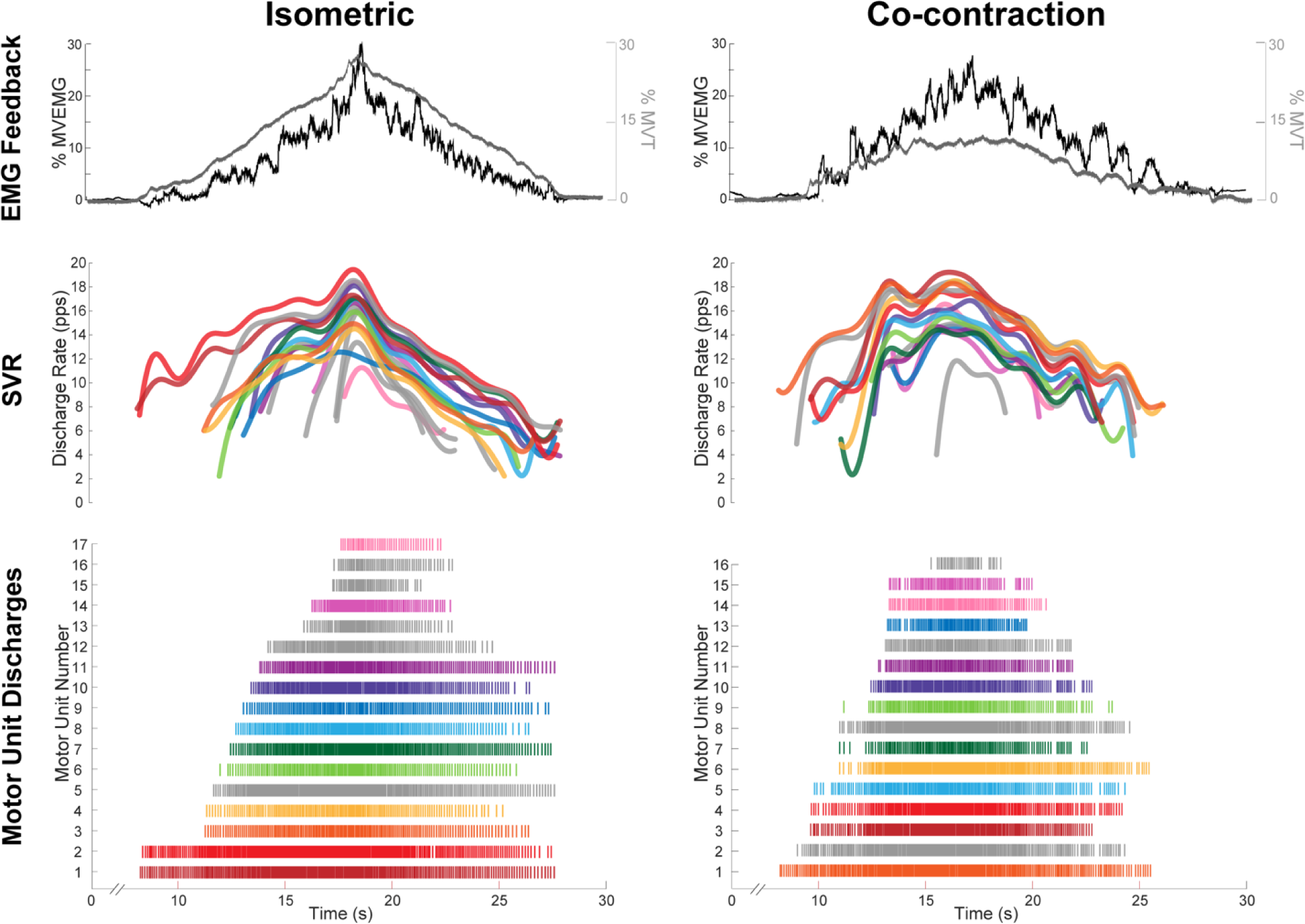
– Example of a single isometric and co-contraction trial from one participant. EMG feedback (black) and dorsiflexion torque (gray) are shown on the top trace. Smoothed motor units with a support vector regression (SVR) are shown in the middle. At the bottom are raster plots showing the individual discharge patterns of units. Units that were matched between the two trials are color coded, and those with no matches are shown in grey.

**Table 1.** Motor unit discharge rates during triangular ramp contractions. Estimated marginal means and standard errors are shown for isometric and co-contraction conditions, with all motor units on the left and tracked motor units on the right. Abbreviations: pps; pulses per second; SE; standard error. Significance noted as: ‘***’ *P* < 0.001, ‘**’ *P* < 0.01, ‘*’ *P* < 0.05.

**A. DISCHARGE RATE AT RECRUITMENT**
|  | *All Data* |  | *Tracked Data* |  |
| --- | --- | --- | --- | --- |
|  | *Mean (pps)* | *SE* | *Mean (pps)* | *SE* |
| **ISOMETRIC** | 9.35*** | 0.390 | 8.62 | 0.40 |
| **CO-CONTRACTION** | 8.75*** | 0.391 | 8.26 | 0.40 |

**B. PEAK DISCHARGE RATE**
|  | *All Data* |  | *Tracked Data* |  |
| --- | --- | --- | --- | --- |
|  | *Mean (pps)* | *SE* | *Mean (pps)* | *SE* |
| **ISOMETRIC** | 15.70*** | 0.668 | 16.9 | 0.52 |
| **CO-CONTRACTION** | 16.00*** | 0.667 | 17.2 | 0.52 |

**C. DISCHARGE RATE AT DERECRUITMENT**
|  | *All Data* |  | *Tracked Data* |  |
| --- | --- | --- | --- | --- |
|  | *Mean (pps)* | *SE* | *Mean (pps)* | *SE* |
| **ISOMETRIC** | 6.84* | 0.243 | 6.41* | 0.314 |
| **CO-CONTRACTION** | 7.08* | 0.243 | 6.97* | 0.314 |

### Motor Unit Discharge Duration Differs between Conditions

To gain additional insight about the timing of discharge in relation to the triangular target that participants were given, we quantified the duration of motor unit discharge (Table 2). In the full dataset ascending phase duration was higher in isometric contractions (χ^2^ = 11.457, *P* < 0.001, *d* = 0.246). Descending phase duration was higher during co-contraction (χ^2^ = 7.8472, *P* = 0.005, *d* = −0.163) and the ascending-descending phase ratio was larger during co-contraction (χ^2^ = 17.026, *P* < 0.001, *d* = 0.24). For tracked MUs, ascending phase duration was higher during isometric (χ^2^ = 5.908, *P* < 0.015, *d* = 0.296), but descending phase duration was similar between conditions (χ^2^ = 1.792, *P* < 0.181, *d* = −0.164). The ascending-descending phase ratio was also similar between conditions in the tracked data (χ^2^ = 1.107, *P* = 0.293, *d* = 0.128).

**Table 2.** Motor unit discharge rates duration during triangular ramp contractions. Estimated marginal means and standard errors are shown for isometric and co-contraction conditions, with all motor units on the left and tracked motor units on the right. Abbreviations: sec; seconds; SE; standard error. Significance noted as: ‘***’ *P* < 0.001, ‘**’ *P* < 0.01, ‘*’ *P* < 0.05.

**A. ASCENDING PHASE DURATION**
|  | All Data |  | Tracked Data |  |
| --- | --- | --- | --- | --- |
|  | Mean (sec) | SE | Mean (sec) | SE |
| **ISOMETRIC** | 5.61*** | 0.219 | 6.15* | 0.201 |
| **CO-CONTRACTION** | 5.34*** | 0.218 | 5.82* | 0.202 |

**B. DESCENDING PHASE DURATION**
|  | All Data |  | Tracked Data |  |
| --- | --- | --- | --- | --- |
|  | Mean (sec) | SE | Mean (sec) | SE |
| **ISOMETRIC** | 6.86** | 0.235 | 6.93 | 0.251 |
| **CO-CONTRACTION** | 7.15** | 0.234 | 7.11 | 0.251 |

**C. ASCENDING-DESCENDING PHASE RATIO**
|  | All Data |  | Tracked Data |  |
| --- | --- | --- | --- | --- |
|  | Mean (sec) | SE | Mean (sec) | SE |
| **ISOMETRIC** | -0.121*** | 0.0308 | -0.102 | 0.029 |
| **CO-CONTRACTION** | -0.174*** | 0.0305 | -0.124 | 0.029 |

To identify a mechanism responsible for different discharge patterns in the two conditions we estimated PIC magnitude. Estimates of PICs (ΔF) were higher in the isometric condition, in both the full dataset (χ^2^ = 72.663, *P* < 0.001, *d* = 0.641) and the tracked units (χ^2^ = 14.597, *P* < 0.001, *d =* 0.656; Figure 5A).

**Figure 5.**
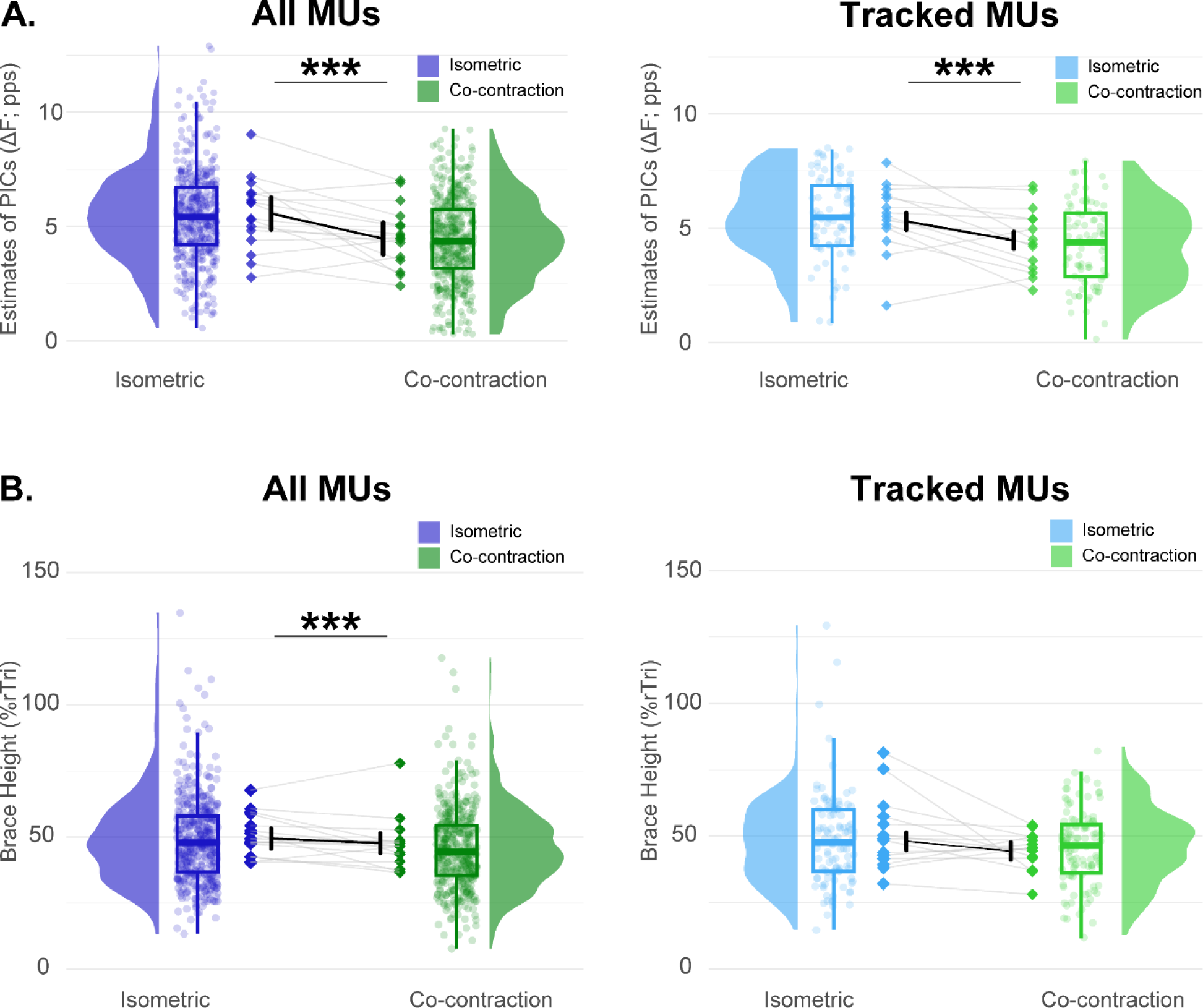
– Estimates of persistent inward currents (top) and brace height (bottom) from the tibialis anterior during isometric (blue) and co-contraction (green). The entire dataset is shown to the right of each plot and the tracked dataset is shown to the left. Model estimates are shown in the dark black lines, individual participant means are shown as diamonds. Box plots show the 25^th^, 50^th^(median), and 75^th^ quartiles, with whiskers showing the 1.5 interquartile range. Distributions across all participants of the respective measure are shown. Significance noted as: ‘***’ *P* < 0.001, ‘**’ *P* < 0.01, ‘*’ *P* < 0.05.

To further probe potential differences in the pattern of inhibition and neuromodulatory to the motoneuron pool during the two tasks, we utilized a quasi-geometric analysis of the individual motor unit discharge rate profiles with respect to the EMG feedback that participants received. Brace height was higher in the isometric condition than co-contraction for the full data (χ^2^ = 19.655, *P* < 0.001, *d* = 0.254) and, although the model did not indicate significant differences, trended higher in the tracked dataset (χ^2^ = 3.8287, *P* = 0.0503, *d =* 0.283; Figure 5A). The acceleration and attenuation slopes did not differ between conditions in both the full (ACC χ^2^ = 2.73, *P* = 0.098, *d =* 0.126; ATT χ^2^ = 0.226, *P* = 0.634, *d* = 0.034) and tracked (ACC χ^2^ = 0.195, *P* = 0.659, *d* = −0. 0612; ATT χ^2^ = 0.454, *P* = 0.501, *d = -*0.098) datasets. The angle between the acceleration and attenuation phases was greater during isometric than co-contraction (ANG χ^2^ = 18.117, *P* < 0.001, *d* = 0.34) but similar between conditions in the tracked units (ANG χ^2^ = 0.151, *P* = 0.698, *d* = 0.056).

Finally, to gain insight about whether there were shifts in the recruitment threshold of motor units, we assessed the amplitude of EMG where the motor unit was recruited. Motor unit recruitment EMG was similar between conditions in the full (χ^2^ = 0.0008, *P* = 0.978, *d* = −0.003) but was higher for isometric contractions of the tracked datasets (χ^2^ = 11.228, *P* = 0.001, *d* = 0.412; Table 3). This indicates that the recruitment thresholds (in terms of agonist EMG) were reduced during co-contraction.

**Table 3.** Geometric metrics and recruitment electromyogram amplitude during triangular ramp contractions. Estimated marginal means and standard errors are shown for isometric and co-contraction conditions, with all motor units on the left and tracked motor units on the right. Abbreviations: pps; pulses per second; SE; standard error; Tri; triangle; **%MEMG; percentage maximum EMG. Significance noted as: ‘***’ *P* < 0.001, ‘**’ *P* < 0.01, ‘*’ *P* <0.05.**

| **A. ACCELERATION SLOPE** |  |  |  |  |
| --- | --- | --- | --- | --- |
|  | *All Data* |  | *Tracked Data* |  |
|  | *Mean (pps/*<br>*%MEMG)* | *SE* | *Mean (pps/*<br>*%MEMG)* | *SE* |
| **ISOMETRIC** | 1.41 | 0.078 | 1.55 | 0.227 |
| **CO-CONTRACTION** | 1.25 | 0.079 | 1.68 | 0.222 |
| **B. ATTENUATION SLOPE** |  |  |  |  |
|  | *All Data* |  | *Tracked Data* |  |

|  | *Mean (pps/*<br>*%MEMG)* | *SE* | *Mean (pps/*<br>*%MEMG)* | *SE* |
| --- | --- | --- | --- | --- |
| **ISOMETRIC** | 0.252 | 0.0184 | 0.267 | 0.0358 |
| **CO-CONTRACTION** | 0.245 | 0.0180 | 0.289 | 0.0356 |

**C. ANGLE**
|  | *All Data* |  | *Tracked Data* |  |
| --- | --- | --- | --- | --- |
|  | *Mean*<br>*(degrees)* | *SE* | *Mean*<br>*(degrees)* | *SE* |
| **ISOMETRIC** | 237*** | 2.96 | 231 | 3.07 |
| **CO-CONTRACTION** | 229*** | 2.95 | 230 | 3.05 |

**D. MOTOR UNIT RECRUITMENT EMG**
|  | *All Data* |  | *Tracked Data* |  |
| --- | --- | --- | --- | --- |
|  | *Mean*<br>*(%MEMG)* | *SE* | *Mean*<br>*(%MEMG)* | *SE* |
| **ISOMETRIC** | 9.33 | 0.405 | 8.68*** | 0.501 |
| **CO-CONTRACTION** | 9.35 | 0.408 | 7.57*** | 0.501 |

## DISCUSSION

The purpose of this study was to investigate the effects of intentional muscle co-contraction on TA motor unit discharge patterns during submaximal isometric ramp contractions in the lower limb. Our results revealed that voluntary antagonist co-contraction affected motor unit discharge patterns, which reduced estimates of persistent inward currents (ΔF). The novel findings from this study add to our basic understanding of how the interplay between excitatory, inhibitory, and neuromodulatory motor commands shape motor unit discharge rate profiles.

### EMG and torque during isometric and co-contraction

Voluntary co-contraction is a complex task due to the competing activation of antagonist pairs that likely impart inhibition onto each other (i.e., reciprocal inhibition) (Crone *et al*., 1987; Hirabayashi *et al*., 2019). Successful execution requires highly complex commands to compensate for the additional inhibition that is not present during traditional contraction (i.e., only involving the agonist muscle). Increasing and decreasing the intensity of co-contraction, as done in the triangular ramp used in the present study, made the task even more challenging. In fact, four participants were excluded from the sample because of their inability to perform the co-contraction ramp with linear increases and decreases in EMG amplitude without increasing torque about the ankle. Among these participants, some contracted one muscle (e.g., TA) more than the other muscle in the antagonist pair (e.g., SOL), resulting in torque generation at the ankle. Meanwhile, others were unable to gradually contract both sets of antagonist muscles to reach the target intensity. Sixteen participants performed the co-contraction ramp appropriately, as evidenced by similar TA EMG, greater soleus EMG, and minimal torque compared to the dorsiflexion isometric condition (Figure 2).

### Motor unit discharge rates differ between conditions

While the TA EMG activity was similar between conditions (i.e., dorsiflexion and co-contraction), the motor unit discharge characteristics differed. It is well established that the volitional activation of an agonist muscle generates Ia afferent input that, through intraspinal circuits, inhibits its antagonist pair (i.e., reciprocal inhibition) (Crone *et al*., 1987; Hirabayashi *et al*., 2019). Therefore, it is reasonable to consider that during co-contraction the simultaneous contraction of antagonist muscles (i.e., plantarflexor muscles) would induce inhibition that would interfere with agonist motor unit discharge (i.e., TA muscle). During co-contraction, the motor unit discharge rate at recruitment was lower, while the peak discharge rate and the discharge rate at derecruitment were higher compared to isometric dorsiflexion. Previous studies that applied vibratory stimuli to the plantarflexor muscles also observed alterations in TA discharge rate characteristics and suggested that these changes were induced by the reciprocal inhibition mechanism (Mesquita *et al*., 2022; Pearcey *et al*., 2022; Orssatto *et al*., 2022). Even though we acknowledge that reciprocal inhibition is a factor influencing motor unit discharge rates, the changes we observed diverge from those reported in previous studies. Here we observed a reduction in the discharge rate at recruitment in the condition that induces reciprocal inhibition (i.e., co-contraction), whereas previously we (Pearcey *et al*., 2022) observed an increase with antagonist vibration. Similarly, while we observed an increase in peak discharge rate and discharge rate at derecruitment in the condition affected by reciprocal inhibition (i.e., co-contraction), previous studies reported an increase in discharge rate when reciprocal inhibition was present (Mesquita *et al*., 2022; Pearcey *et al*., 2022; Orssatto *et al*., 2022). Reciprocal inhibition in these previous studies was induced by antagonist nerve stimulation (Mesquita *et al*., 2022) or antagonist tendon vibration (Pearcey *et al*., 2022; Orssatto *et al*., 2022), whereas currently reciprocal inhibition was induced by the voluntary co-contraction of the antagonist muscle. It has been demonstrated that contraction of remote muscle groups can amplify motoneuron excitability, probably through increased monoamines delivered by descending tracts from the brainstem (Heckman *et al*., 2008; Wei *et al*., 2014; Orssatto *et al*., 2022). Thus, the voluntary co-contraction of the antagonist muscle may have not only caused reciprocal inhibition but also increased the excitatory and neuromodulatory commands to the motoneurons.

### Estimates of PICs were lower during co-contraction

Since PICs are highly sensitive to inhibition, reciprocal inhibition from contraction of antagonist pairs could reduce PICs in the agonist muscle. However, contracting remote muscle groups can increase PICs (Heckman *et al*., 2008; Wei *et al*., 2014; Orssatto *et al*., 2022), and thus antagonist muscle contraction could increase monoaminergic drive and facilitate PICs during the co-contraction task. Here, we analyzed the combined effect of these two potential alterations in motor commands and find that local inhibition from antagonist muscle contraction appears to overcome any potential increase in neuromodulatory drive.

Our findings corroborate previous studies that have also observed a PIC reduction in the presence of reciprocal (Mesquita *et al*., 2022; Pearcey *et al*., 2022; Orssatto *et al*., 2022). As mentioned above, these previous studies relied on methods to induce sensory input rather than direct activation of the antagonist muscle (Mesquita *et al*., 2022; Pearcey *et al*., 2022; Orssatto *et al*., 2022). It is probable that these methods do not elicit the same monoaminergic input as the voluntary contraction of the antagonist muscle, which was specifically investigated in our study. In this regard, our results complement the findings of Orssatto et. al. (2022) showing that the reciprocal inhibition input overlaps the increased monoaminergic drive triggered by the voluntary activation of the antagonist muscle. Furthermore, vibration applied to the antagonistic muscle can activate additional sensory inputs, such as non-local mechanoreceptors or heteronomous muscle spindles, which can influence the excitability of motoneurons (Garnett & Stephens, 1981; Barss *et al*., 2021). Nevertheless, our results revealed that reciprocal inhibition induced by voluntary antagonist contraction elicited a similar reduction in ΔF (i.e. 1.1 pps, 19.7%) to those observed in the previous studies (Pearcey *et al*., 2022; Orssatto *et al*., 2022) that used vibratory stimuli (0.54 pps, 14.4%; 0.72 pps, 14.7%, respectively).

In summary, our results demonstrate that voluntary co-contraction represents an intriguing paradigm to facilitate the concurrent assessment of excitatory, neuromodulatory, and inhibitory inputs. Co-contraction elicits reciprocal inhibition and increased neuromodulatory input, which results in decreased PICs and alterations in the discharge rate profiles of motor units. Given that co-contraction has been employed as a resistance training method, forthcoming studies could explore whether prolonged co-contraction training induces enduring adaptations in the intrinsic properties of motor units.

## METHODOLOGICAL CONSIDERATIONS

Due to the difficulty in performing the task, we assessed only one submaximal level of contraction (i.e., 30% MVC). Since contraction intensity has profound effects on the composition of motor commands (Škarabot *et al*., 2023) future investigations with higher intensity co-contractions are needed to verify whether the reciprocal inhibition input will continue to overcome the increased monoaminergic drive resulting from the voluntary activation of the antagonist muscle. Another limitation is that we only analyzed the isometric condition in a single ankle position (i.e., at 100°). Alterations in the ankle angle can modify passive and active joint moments (Jamwal *et al*., 2017), contractile properties of the muscles (i.e., tension-length relationship) (Rassier *et al*., 1999; Cadeo *et al*., 2023) and, in particular, motor unit discharge rate profiles and estimates of PICs (Beauchamp *et al*., 2023*b*). Thus, a different ankle position could influence the recruitment threshold and discharge rate of the motor units required to maintain zero torque at the ankle during voluntary co-contraction. We also only assessed estimates of PICs and discharge rate patterns from the TA. It is important to consider that various muscles have diverse innervation (Banks, 2006; Kissane *et al*., 2023), which may affect the extent of reciprocal inhibition. Finally, there were discrepancies in our findings when analyzing all motor units compared to analyzing only tracked motor units. This likely is due to a lower sample size of tracked motor units across tasks, or could be due to a difference in recruitment threshold between the two conditions of the tracked motor units.

## PRACTICAL APPLICATIONS

In the last decade, voluntary co-contraction has been proposed as a method of strength training. To do this, a person perform sets of voluntary simultaneous contraction (i.e., co-contraction) of antagonistic pairs (e. g. elbow flexors and extensors) with no external apparatuses for loading (Mackenzie *et al*., 2010; Zbidi *et al*., 2017; Fujita *et al*., 2021). Previous studies have indicated that co-contraction training promotes strength gain (Mackenzie *et al*., 2010; Villalba *et al*., 2024) and hypertrophy (Counts *et al*., 2016), similar to conventional resistance training, which makes this method very promising with potential application in microgravity and rehabilitation backgrounds. It is now apparent that the neural commands required to perform co-contraction differ from those to perform agonist contractions. These novel insights are likely to shed light on the application of these types of muscle contraction for adaptations in both health and disease.

## CONCLUSION

Voluntary antagonist co-contraction significantly altered motor unit discharge characteristics and reduced estimates of persistent inward currents. The novelty of our approach, which concurrently considers both inhibitory and excitatory inputs arising from voluntary antagonist co-contraction, enhances our basic understanding of the interplay between excitatory, inhibitory, and neuromodulatory motor commands that shape motor unit discharge rate profiles. These findings also hold promise for optimizing therapeutic strategies and training protocols that utilize voluntary muscle co-contraction.

### Data availability statement

The data that support the findings of this study are available on request from the corresponding author.

### Author contributions

M.M.G., S.T.J., J.A.B., C.J.H., and G.E.P.P. conceptualized and designed the research; M.M.G., S.T.J., J.A.B., and G.E.P.P performed the experiments; F.N developed blind source separation algorithms; M.M.G., S.T.J., and J.A.B. analyzed the data; M.M.G., S.T.J., J.A.B., C.J.H., and G.E.P.P. interpreted the results of experiments; M.M.G., S.T.J., and J.A.B. prepared the figures; M.M.G. and S.T.J. drafted the manuscript; M.M.G., S.T.J., J.A.B., F.N., C.J.H., and G.E.P.P. revised and approved the final version of the manuscript.

### Funding

This work was supported by the São Paulo Research Foundation (FAPESP; grant #2020/03282- 0), the National Institute of Health (NIH; grants R01NS098509−01, R01NS098509−05, and NINDS F31 NS120500), and the Natural Sciences and Engineering Research Council of Canada (NSERC; Discovery Grant 2023−05862, and Discovery Launch Supplement 2023−00279).

